# Signal separability in integrated neurophotonics

**DOI:** 10.1101/2020.09.27.315556

**Authors:** Dimitri Yatsenko, Laurent C. Moreaux, Jaebin Choi, Andreas S. Tolias, Kenneth L. Shepard, Michael L. Roukes

## Abstract

A new modality Photonic probes record fluorescent signals by using arrays of light emitters and detectors embedded in neural tissue. Neither the emitted nor collected light fields are focused. Instead, in proposed configurations, hundreds of emitters will form rapid sequences of structured illumination patterns—providing sufficient spatial and temporal differentiation of neural signals for computational demixing. Here we define criteria for evaluating probe designs for achieving better signal separability. We find that probe geometry has profound, often unintuitive, effects on the separability of neural signals, providing initial design guidelines to achieve separation of individual cells in densely labeled populations.

## Introduction

Integrated neurophotonics is a proposed neurophysiology modality for large-scale recordings and stimulation of neural circuits *in vivo* [1]. Neurophotonic probes comprise multiple shanks that carry arrays a light emitting elements (E-eixels) and light detecting elements (D-pixels). Inserted into brain tissue, they produce rapid sequence of structured illumination patterns, allowing to resolve and localize optical signals from voltage- or calcium-sensing fluorescent proteins.

The **central question** addressed in this paper is: How does the design of a photonic probe affect its ability to resolve the optical signals from populations of fluorescently labeled neurons in the embedding tissue.

Fluorescence imaging modalities are distinguished by how they structure their illumination and how they focus the collected light (Figure 1). Wide field microscopy, for example, uses a wide illumination field and focuses the collected light optically, thereby separating signals by the spatial location (Fig. 1A). The entire field can be imaged simultaneously. In laser-scanning microscopy, the roles of light illumination and collection are revered: the illumination light is focused and the collected light is detected without discrimination (Figure 1B). Since the collected photons are not sorted, focused illumination must be structured in time by, for example, raster scanning. New capabilities can be gained by focusing both the illumination and collection fields as in, for example, confocal imaging and lightsheet microscopy [2].

**Figure 1:**
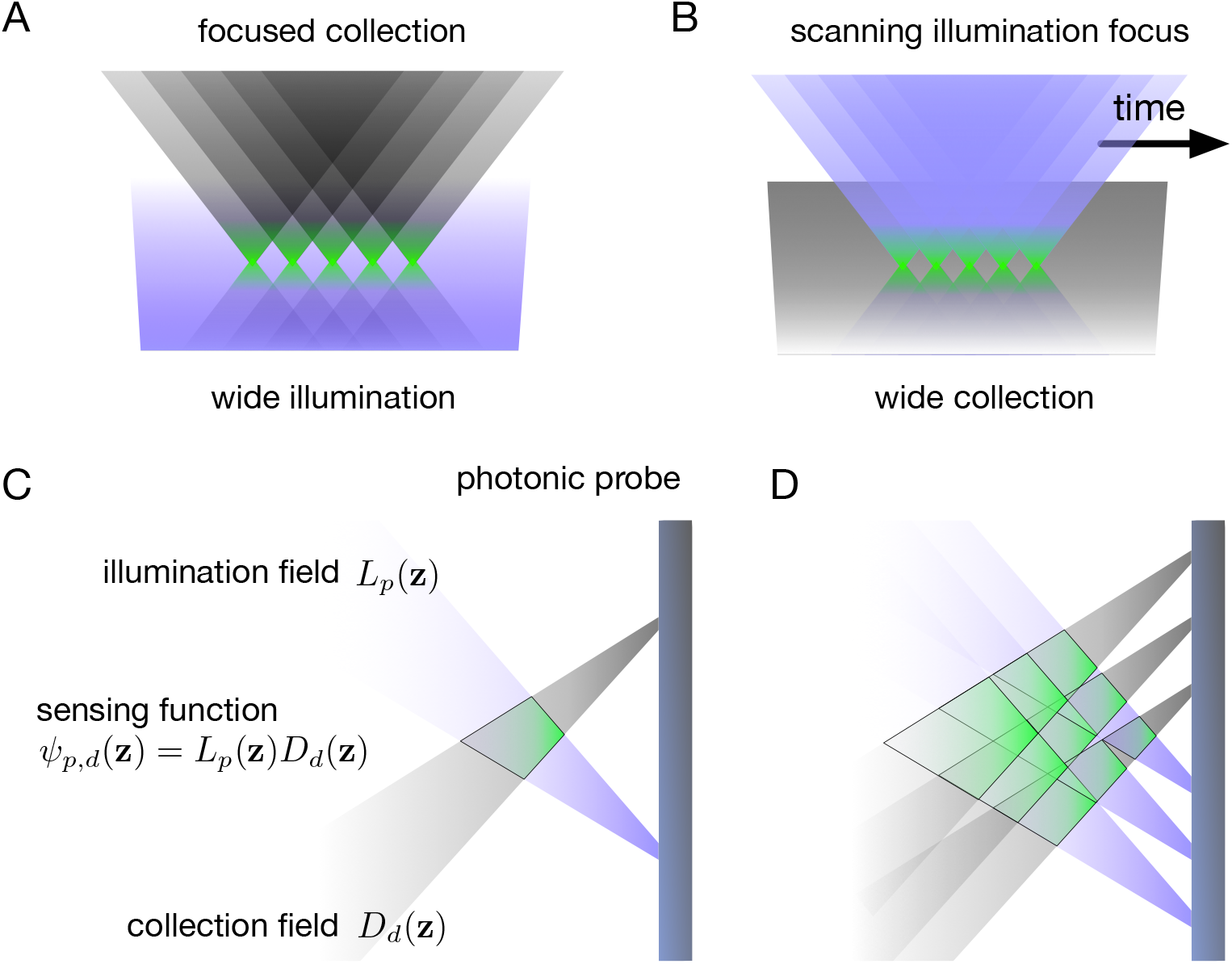
The structures of the collection and illumination fields in various types of fluorescence imaging. A: In wide-field microscopy, the illumination field is wide while the collection fields are focused. B: In laser-scanning microscopy, the collection field is wide while the illumination field is focused on one point at a time. C: In photonic probes, neither the illumination nor the collection fields are focused but have fine spatial structure thanks to being embedded in tissue. Their overlap (pointwise product) forms the illumination-collection function (ICF). D: The number of ICFs is the product of the number of illumination fields and the number of overlapping detection fields.

In contrast, neurophotonic probes are lensless and do not focus either the illumination nor the illumination fields (Figure 1C and D). For any given E-pixel/D-pixel pair, the pointwise product of the E-pixel’s illumination field and the D-pixel’s collection field form their *sensing function*, or the illumination-collection function (ICF). Furthermore, the illumination fields emitted by E-pixels may be spatial modulated or steered by changing the light’s wavelength, further multiplying the number of possible sensing functions. In practical acquisition protocols, multiple E-pixels will activate simultaneously to form complex illumination fields. Generally, the maximum number of linearly independent ICFs is the product of the number of illumination fields and the number of detection fields used in the acquisition. The probe acquires signals by rapidly cycling through the entire set of illumination patterns, with the entire cycle taking a period on the scale of a millisecond. During each pattern, all detectors record a photon count. Each ICF corresponds to a recorded *channel*, a single measured photon count in given sample comprising the full illumination cycle.

The entire ICF ensemble characterizes the optical resolving power of the photonic probe. ICFs can be thought of as a generalization of the point-spread-function (PSF) as the description of optical performance of a conventional imaging system where single PSF describes the optical resolution anywhere across the entire image. In contrast, ICFs are non-stationary: they are much larger, irregular, diverse in their shapes and intensities. They substantially overlap and are bound to their specific locations. Furthermore, the ICFs depend on the optical properties of the imaged volume and cannot be assumed to be known precisely and may even change slightly over time due to tissue movements, blood vessel constriction/dilation, and changes in blood oxygenation.

With no focusing optics, ICFs will generally be too coarse to resolve individual neuronal bodies: each recorded channel will comprise signals from tends or hundreds of cells covered by its ICF. Yet the ICFs can be shaped in such a way that any two neurons, even when only microns apart, will project differently onto the several overlapping ICFs at their locations to produce linearly separable signals. Thus signals acquired by photonic probes must rely on computational analysis for signal separation and localization.

Neuropil signals, *i.e*. signals from myriad other cells’ neurites passing through the imaged volume will contaminate signals, limiting the ability to isolate individual neuronal signals. Photonic probes will rely on the ongoing efforts to constrain fluorescent reporters to the cells’ somatic compartments, thereby reducing the density of coherent signals in the recorded volume [3–5].

The optimal architectures and configurations for recording neuronal signals have not yet been defined. These must be constrained by considerations of their manufacturability, implantability, and permissible power and recording rates. Many aspects of the design present unintuitive tradeoffs. For example, should the D-pixels maximize their sensitivity to collect the most photons or increase their angular selectivity to refine the structure of the probe’s ICFs?

This study defines a framework for a systematic evaluation and optimization of photonic probe designs for neuronal recordings and illustrates a number of important design tradeoffs.

## Results

### Geometric designs

We evaluated three probe geometries: Designs A, B, and C as illustrated in Figure 2.

**Figure 2:**
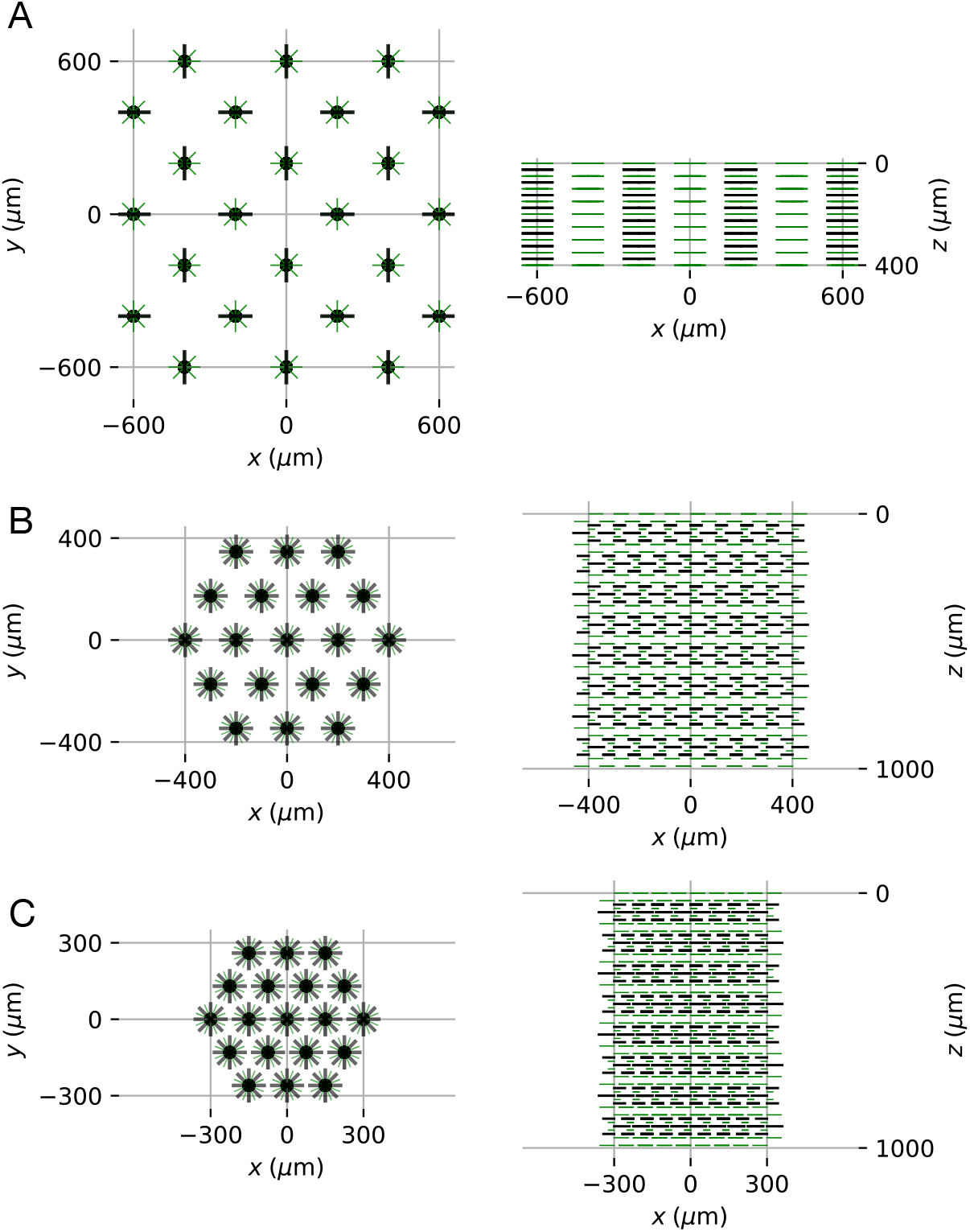
Probe designs A, B, and C. Top view is on the left and side view is on the right. The black dots indicate the shanks; the green ticks indicate the orientation of E-pixels and the dark ticks indicate the orientation of the D-pixels.

Design A comprises 24 shanks, forming a lattice of squares with 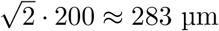 sides (Figure 2A). Each of these shanks supports 72 E-pixels arranged as nine separate rings, each comprising eight E-pixels. Along the length of the shank (z-axis) and between these rings of E-pixels are opposite-facing pairs of D-pixels. In total, the design features 1728 E-pixels and 384 D-pixels. The E-pixels measure 10 μm × 10 μm in size whereas the D-pixels measure 10 μm × 50 μm. The rings of E-pixels and the pairs of D-pixels are arranged uniformly along the length of the shank so that the distance from the center of the top E-pixels to the center of the bottom E-pixel is 400 μm. Adding the height of the E-pixels, the overall active length of shank is 410 μm, which corresponds to the volume of 0.0328 mm^3^ per shank or 0.7872 mm^3^ per probe if these modules are combined as a continuous tiling.

Design B comprises 19 shanks forming a triangular lattice with 200 μm pitch (Figure 2B). Each shank carries 34 E-pixels and 33 D-pixels spaced uniformly along the full 1-mm active length (or 1010 μm accounting for the height of the top and bottom E-pixels). The E-pixels measure 10 μm × 10 μin size, and the D-pixels measure 10 μm × 20 μm. Each pixel’s field is oriented 112.5° clockwise from the pixel immediately above it on the shank, to form a helical pattern. With the total 627 D-pixels and 646 E-pixels, the probe’s working volume is 0.658 mm^3^ (0.0346 mm^3^ per shank).

Design C is identical to B except the pitch is reduced to 150 μm while keeping the same shank length (Figure 2C). The probe’s effective working volume becomes 0.370 mm^3^ (0.0195 mm^3^ per shank).

### Field shaping

In addition to the probe geometry, the shape of the illumination fields of E-pixels and detection fields of D-pixels will affect the properties of acquired signals.

A maximally sensitive D-pixel will have a Lambertian spatial profile (Figure 3 top left). Making it spatially selective without focusing optics will decrease its sensitivity but may confer benefits for refining the selectivity of the resulting ICFs. The selectivity of D-pixels was modulated as cos*^k^ α* where *α* is the incidence angle. Thus cos^0^ *α* described a non-selective Lambertian profile and cos^4^ *α* or cos^8^ *α* produced narrower selective profiles. The detection fields for cos^0^ *α*, cos^4^ *α* and cos^8^ *α* are shown in the top panel of Figure 3. Note that, in diffuse illumination, selectivity profiles cos^4^ *α* and cos^8^ *α* have detector sensitivities of 1/3 and 1/5 of the Lambertian profile, respectively.

**Figure 3:**
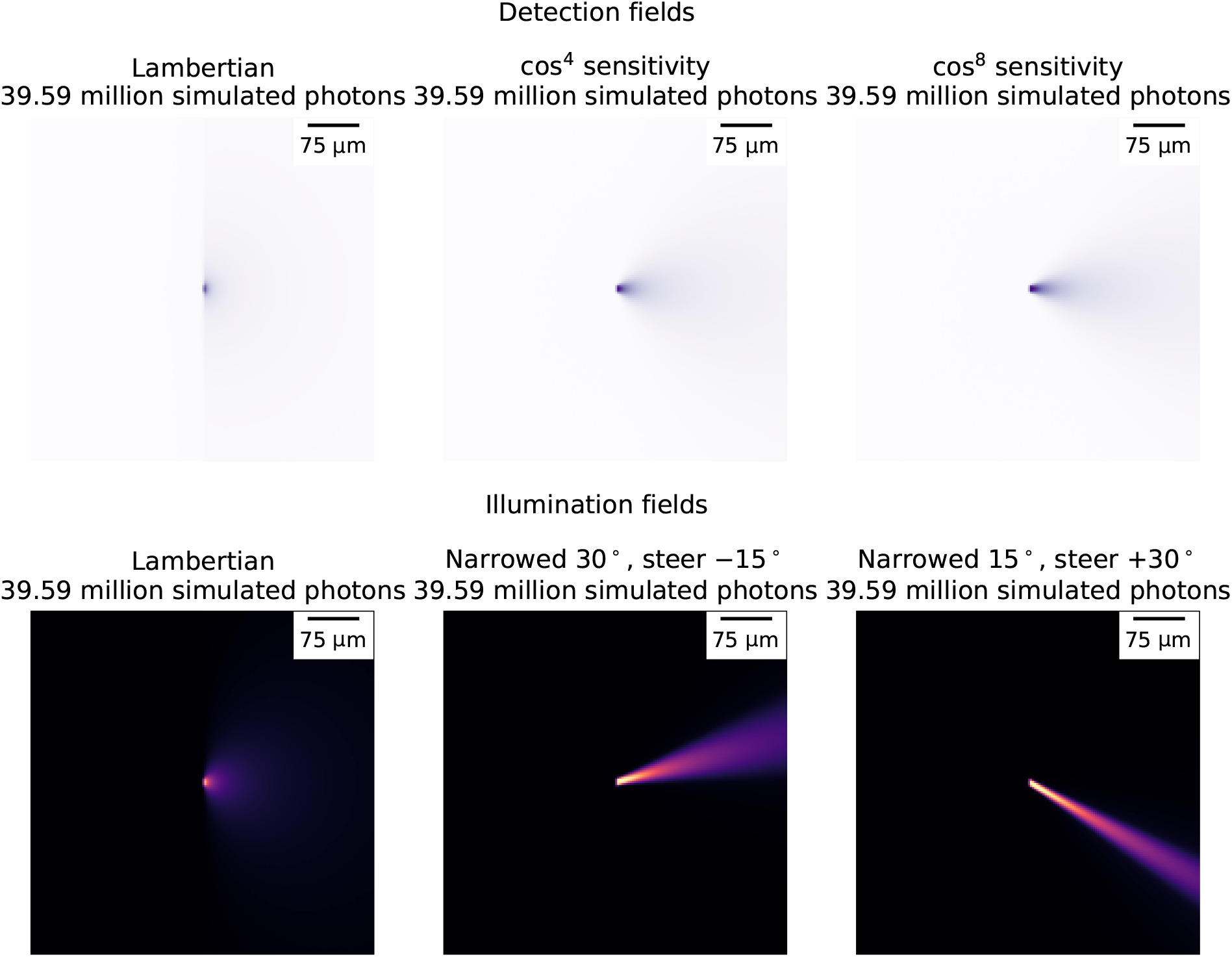
Maximum projections of the simulated detection fields (top row) and illumination fields (bottom row) among those used in the study.

A diffuse emitter produces a Lambertian spatial profile. In our simulated designs, we experimented with compressing the Lambertian profile into a cone of 60°, 30°, and 15°. Furthermore, compressed beams could be steered up and down in the plane of the shank up to 60° to dynamically alter the illumination field between the acquisition frames of the same sample, providing further multiplication of the number and diversity of the probe’s ICFs. All compressed emitter fields were steered at nine discrete angles from −60° to +60° spaced uniformly at 15° increments. Figure 3 bottom row illustrates the illumination fields of a Lambertian (left) and compressed and steered illumination fields (middle, right).

For convenience, we coded the designs to include the geometry, the detector profiles, and the emitter profiles. For example, design A-cos^0^-180° denotes geometry A with Lambertian detector profiles and Lambertian emitter profiles whereas B-cos^8^-15° denotes geometry B with cos^8^ detector profiles, and the emitter fields compressed into 15° cones.

### Separability decreases with labeling density

To evaluate the performance of each geometry, we estimated the signal-to-noise ratios of recorded calcium events due to single action potentials in populations of spiking neurons after demixing the cells’ optical signal from the rest of the population, resulting in “spike SNRs” (See Methods). Here the *signal* is the amplitude of a calcium event due to a single action potential in a given neuron after linear demixing of its activity from from the signals of all the other neurons in the population, after applying a matched filter. And the *noise* is the Poisson-distributed photon shot noise for the same neuron resulting from the quantal nature of the detected light, including the D-pixels’ additive *dark noise*—after applying said demixing and filtering.

Note that the estimation of spike SNRs is possible because the simulation provides direct access to the mixing matrix. In actual experiments, the mixing matrix will not be accessible and the demixing matrix will need to be inferred from the data by applying source separation techniques [6]. This study does not yet propose a specific approach for signal demixing. Rather, we use the known ground truth matrix to evaluate the ability of the design to produce useful signals once the demixing matrix is available.

We calculated the spike SNRs for all cells for several densities of labeled neurons from 1,000 mm^−3^ to 100,000 mm^−3^, under the assumption that the fluorophore molecules were localized in the cells’ somas. Note that 100,000 mm^−3^ approaches the maximum density of neuronal somas in mouse cortex.

Figure 4 depicts the results for design A-cos^0^-180°. The spike SNRs decreased rapidly with increasing labeling density of the neuronal population as the signals from individual cells become washed out by the brighter fluorescence background and also become more mixed with the signals of other cells, requiring more “aggressive” demixing to project out contaminating signals.

**Figure 4:**
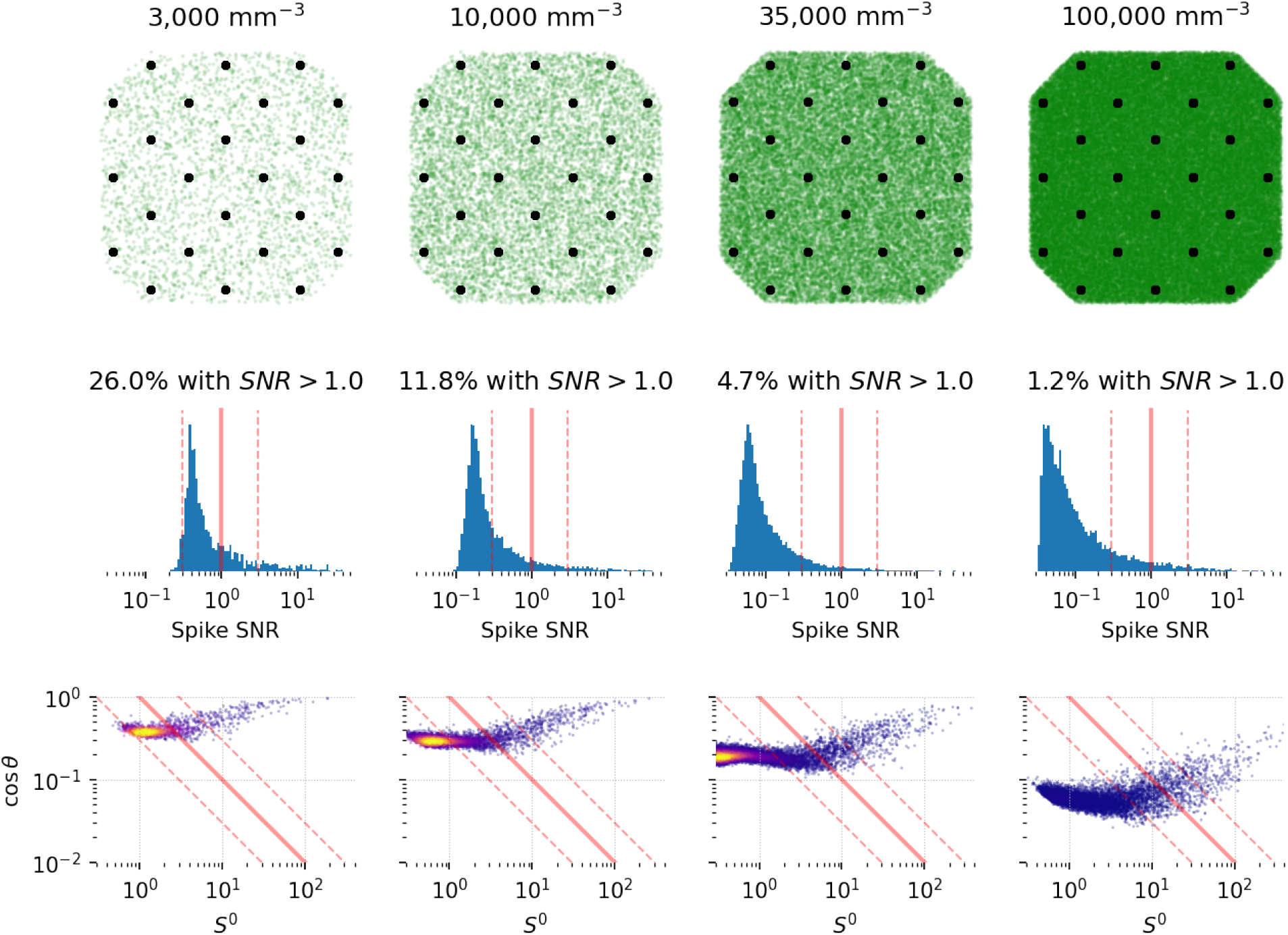
Spike SNRs decrease as the density of labeled cells increases. Top row: a depiction of the neurophotonic probe design A-cos^0^-180° with various densities of labeled neuronal somas from 3,000 to 100,000 mm^−3^. Middle row: historgrams of spike SNRs for each imaging condition; the red lines mark spike SNR levels of 0.3, 1.0 (thick line), and 3.0. Only neurons between the shanks are plotted. Bottom row: the scatter plot of spike SNRs decomposed into the mixed spike SNR *S*^0^ and the cosine of the separation angle cos *θ*. Each dot represents a neuron. The red lines are the same as in the middle row.

We use the cutoff of spike SNR > 1.0 to qualify cells as “separable”—this is an arbitrary but reasonable choice that is employed for comparable current calcium imaging modalities that do not require demixing. At low labeling densities of 3,000 mm^−3^, a fraction (26.0%) of cells met this criterion but this fraction dropped to 11.8% at 10,000 mm^−3^, 4.7% at 35,000 mm^−3^, and only 1.2% at 100,000 mm^−3^. Figure 5 shows these fractions for this and other probe designs.

**Figure 5:**
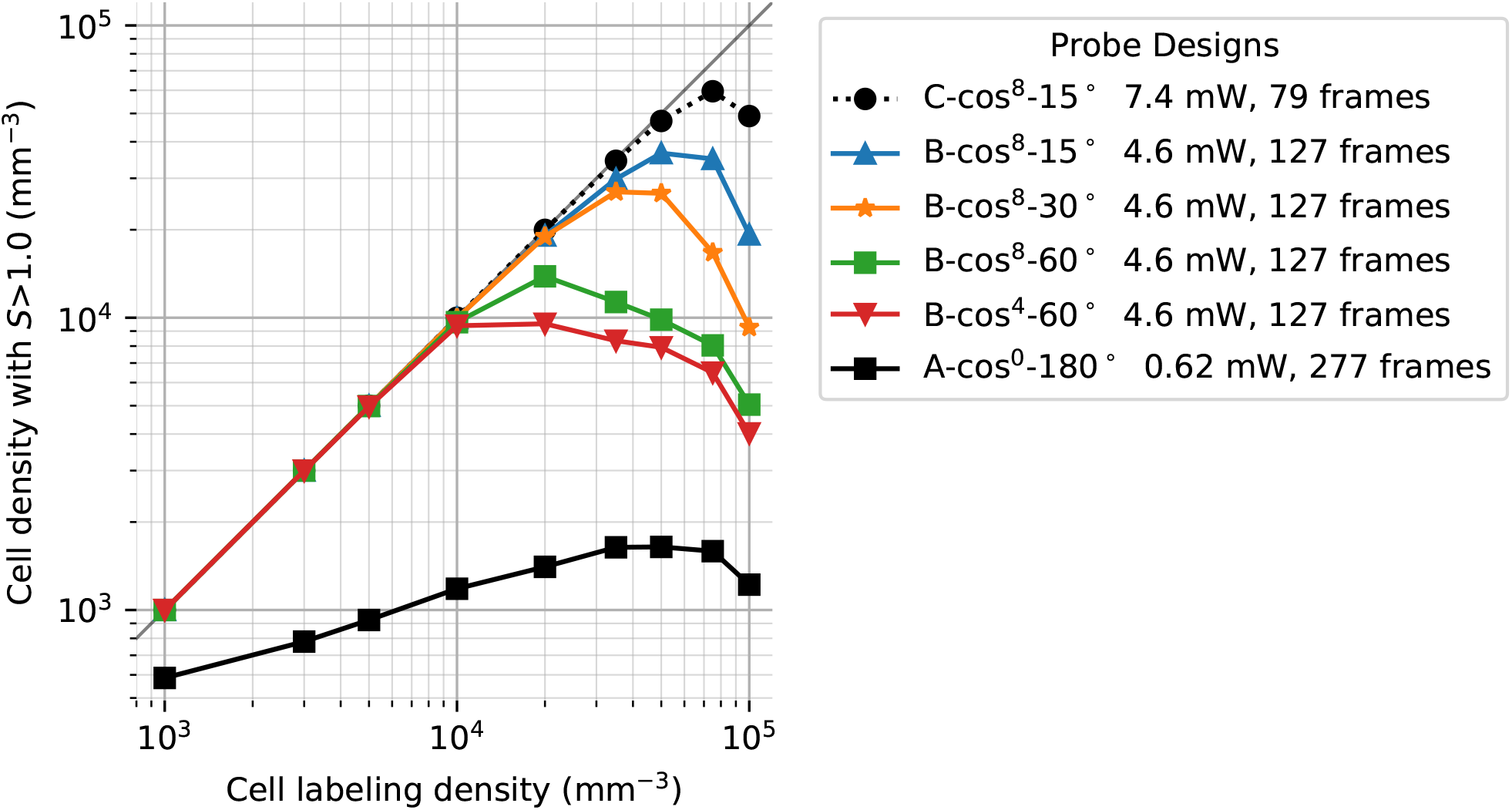
The densities of cells whose demixed spike signal *S* > 1 (Eq. 1), for six different designs, as a function of cell labeling density.

### SNR factorization

To gain insight into the factors limiting spike SNRs, we decomposed them as

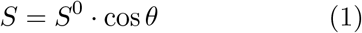

where *S* is the demixed spike SNR of a cell, *S*^0^ is the cell’s *mixed spike SNR, i.e*. the SNR optimally aggregated across all channels under the condition that the fluorescence of all the other cells is known or constant.

The factor cos *θ* expresses the reduction in SNR due to the demixing process required to project out the activities of all the other cells from the signal. Technically, *θ* is the angle between the neuron’s signal and the nearest vector that is normal to the hyperplane formed by the activities of all the other cells in the multidimensional space of the recorded signal (channels=dimensions).

The two components (*S*^0^) and (cos *θ*) may be considered to express the neuronal population’s “exposure” and “focus” within the probe’s recorded volume.

High values of mixed spike SNRs (*S*^0^) indicate that the cells are well positioned in bright illumination by multiple E-fields and their emitted fluorescent light is well detected by multiple D-pixels. The value of *S*^0^ can be increased by improving the detector efficiency and increasing light intensity. It also scales predictably in inverse proportion square root of the labeling density as the background intensity increases proportionally.

The separation cosine, cos *θ*, expresses the spatial resolving power of the probe, which relates to the geometric properties of the probe and its ICFs. Compact, minimally overlapping ICFs will produce nearly orthogonal separation, with cos *θ* approaching 1.0. The values of cos *θ* can drop precipitously with labeling density as cells become less resolvable.

The bottom row of Figure 4 decomposes the Spike SNR into its components for design A-cos^0^-180°. Since this design uses diffuse emission and detection with Lambertian profiles, all but a small fraction of cells have the same low cos *θ* that decreases rapidly with labeling density. A small population of cells closest to the shanks has both high *S*^0^ and cos *θ* with very high SNRs even at highest labeling densities. These cells, numbering in hundreds, will be separable by blind source separation techniques against the background “noise” activity of all other cells.

### Field shaping increases separability

The first design, A-cos^0^-180° demonstrates only moderate performance even at low labeling densities (Figures 4, 5, and 6), even though a small minority of cells close to the shanks produce high spike SNRs even at highest labeling densities. With its 1728 E-pixels and 384 D-pixels, to achieve the high number of channels required for demixing cells, the number of frames in a cycle must be kept sufficiently high. The illumination cycle optimization algorithm settled on 277 frames with only 6.2 E-pixels active at a time, on average, or 624 μW of power. In all the illumination cycle, each E-pixel was turned on only in one frame. Activating E-pixels several times during each sample would increase the exposure but would decrease the distinctions between the illumination patterns. As a result, the design was underpowered in both exposure and resolving power.

**Figure 6:**
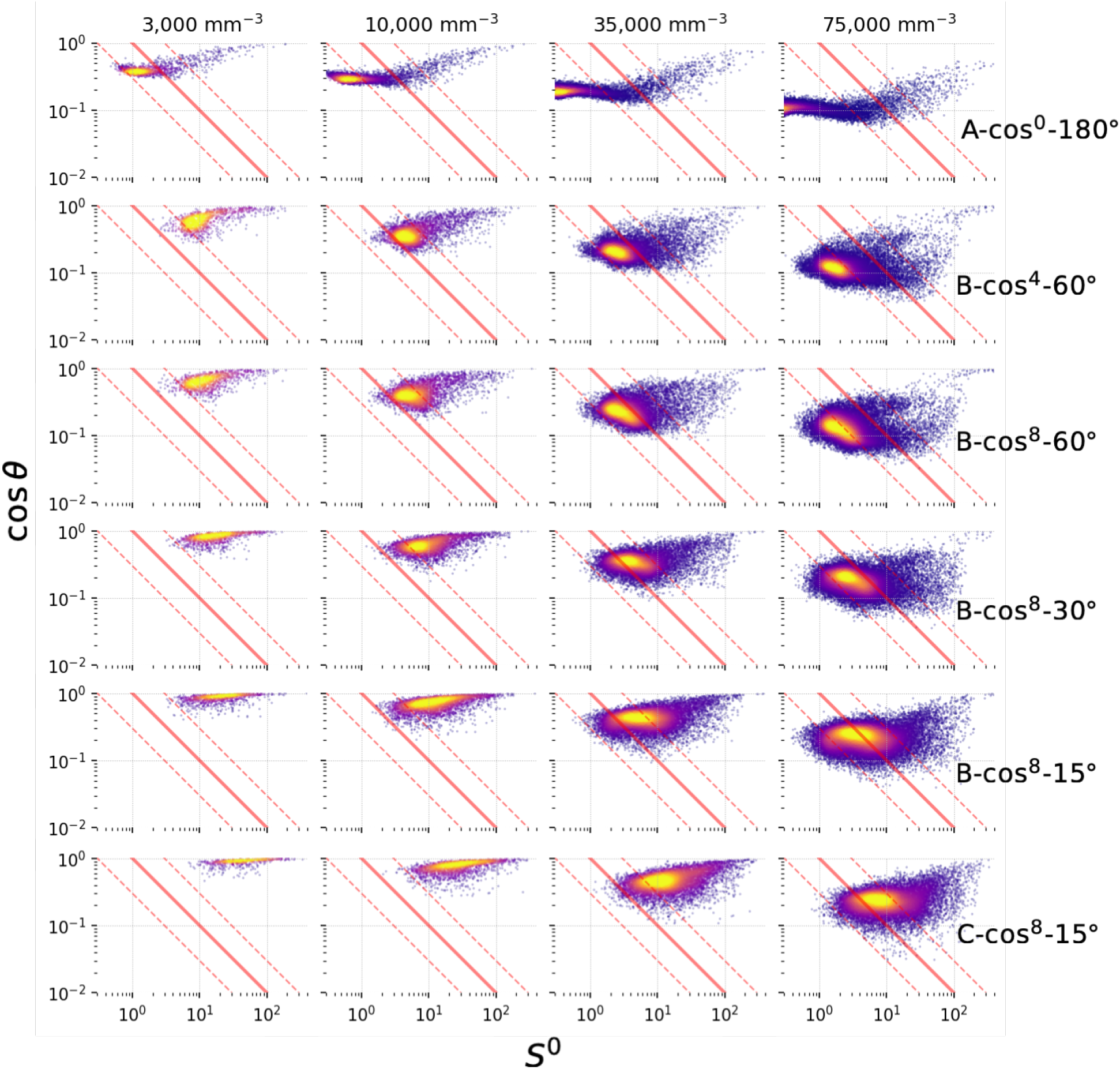
The spike SNRs decompositions as *S* = *S*^0^ · cos *θ*. Each dot represents a neuron. The red lines indicate the isolines *S* = 0.3, *S* = 1.0 (thick), and *S* = 3.0

We experimented with several probe characteristics. First, enabling beam steering in E-pixels allows turning them on in multiple frames of the illumination cycle, increasing average power, with different illumination fields thus increasing the exposure without sacrificing the separation between the illumination fields [**?**, sacher2019beam]

Second, the illumination fields of the E-pixels and D-pixels could be narrowed to refine the structure of the resulting ICFs.

Third, the spacing between the shanks can be reduced.

These adjustments produced a substantial increase in performance (Figures 5 and 6). In Design B, the illumination cycle had 127 frames with 45.8 E-pixels active in each frame on average, resulting in 4.58 mW of power and highly diverse illumination fields.

Remarkably, increasing the selectivity of D-pixels further improved performance. From Design B-cos^4^-60° to Design B-cos^8^-60°, the selectivity was narrowed so that in diffuse light, the efficiency of the D-pixels dropped by 40% but the increased selectivity more than compensated for that loss.

Also narrowing the E-pixels’ spatial profiles substantially increased the performance, *e.g*. from Design B-cos^8^-60° to B-cos^8^-30° and then B-cos^8^-15°, the number of cells above the threshold increased several fold at high labeling densities.

Reducing the spacing between the shanks allows reducing the number of channels, keeping more E-pixels activated in each frame. The illumination cycle for Design C has only 79 frames with 73.6 E-pixels active in each frame on average, resulting in 7.36 mW of power combined with high selectivity, reaching 50% of cells at *S* > 1.0 at 100,000 mm^−3^ labeling density.

## Methods

All simulations were performed using the Data-Joint framework [7]. The complete computational workflow to reproduce the results is published in open source at https://github.com/dimitri-yatsenko/photix.

### Simulated light emission, transport, fluorescence, and detection

Light transport simulation was performed using a Monte-Carlo simulation. The parameters are summarized in Table 1.

- Wavelength λ = 480 nm corresponding to *ν* = 2.4 × 10^18^ photons·J^−1^.
- The same scattering and absorption mean lengths were used for both excitation and emission light: *L_sc_* = 50 μm (scatter) and *L_ab_* = 15,000 μm (absorption). For scatter, the anisotropy coefficient in the Henyey-Greenstein scattering function was set at *g* = 0.88.
- The average power of light emitted by the E-pixels when activated was set at *w_E_* = 100 μW.
- Fluorophores were assumed to be at *F*_0_ = 0.05 of maximum fluorescence on average (including baseline fluorescence and spiking activity).
- The effective cross-section of a neuron was set at *σ* = 0.1 μm^2^ at maximum fluorescence. This was estimated based on 10 μM dye concentration in a spherical volume of 7 μm radius, making 8.6 × 10^6^ molecules per cell and 2.3 × 10^−16^ cm^2^ cross-section of each fluorophore molecule, and the quantum yield of *ϕ* = 0.6 [8].
- A calcium spike due to an action potential was assumed to produce a peak increase of fluorescence of *δ* = 0.02 (Δ*F/F* = 0.4) with exponential decay with time constant *τ*_Ca_ = 1.5 s. These parameters are consistent with state-of-the art calcium indicators [3].
- D-pixels yielded discrete photon counts with quantum efficiency *η* = 0.6 and an additive dark noise level of *ν* = 300 s^−1^, which corresponds to the performance of state-of-the-art single photon avalanche diode (SPAD) detectors [9].
- Importantly, the simulation uses σ · *w_E_* · *F*_0_ · *η* jointly as a product. Therefore these four parameters can be adjusted in inverse proportion without modifying the result of the simulation. In our companion paper [1], the same simulation settings were used but the E-pixel power was reported as *w_E_* = 20 microwatts and the cross-section was increased to 0.5 μm^2^ to compensate.

**Table 1:**
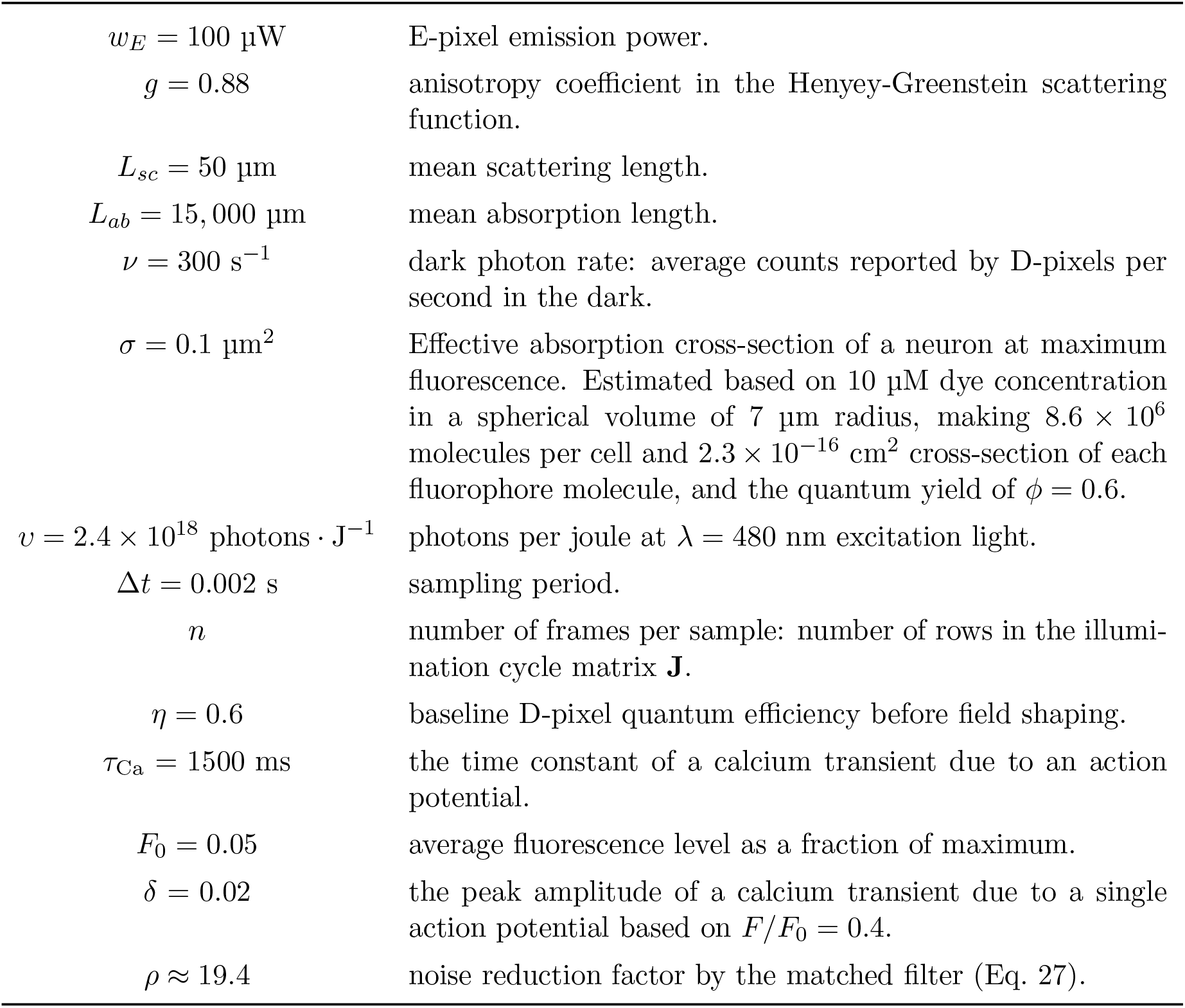
Summary of notation and model parameters.

The light transport simulation yielded the photon detection probability fields *D*(**z**) (unitless) for D-pixels and the relative irradiance field *L*(**z**) (μm^−2^) for E-Pixels.

*D*(**z**) expresses the probability that a photon emitted in a random direction from point **z** will be counted by the given D-pixel.

*L*(**z**) expresses the fraction of photons emitted by the given E-field expected to pass through a sphere with cross-section of 1 μm^2^ at point **z**.

Figure 3 illustrates examples of such fields. We modeled the illumination and collection fields of each shape once and sampled from them to model the individual E- and D-pixels on the shank. This approach provides great computational efficiency but does not allow the particular details specific to each pixel with shank shadows and any modeled tissue. While this approach is sufficient for preliminary optimization, a more detailed simulation will be carried out with modeling each element in its place in tissue. It is not yet clear what effect the lack of shadows has on signal separability since shadows decrease the illumination and detection but they also increase the diversity of ICFs, possibly aiding signal separability.

Populations of neurons were modeled as points in space **z***_i_*, *i* = 1, …, *m* with a minimal pairwise distance of 8 μm at densities between 1,000 and 100,000 mm^−3^ in the volume embedding the probe, including a 75 μm margin surrounding the outer periphery of the multi-shank module as illustrated in Figure 4. The points outside the convex hull of the probe itself were included in the simulation (as contributing to a contaminating fluorescence background) but were excluded from the evaluation of spike SNRs.

### Fluorescence matrix

The fluorescence matrix **F** (E-fields × neurons ↦ probability) comprises elements

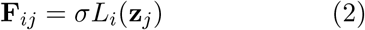

expressing the probability that a photon emitted from by the *i*^th^ E-field will be re-emitted by the *j*^th^ neuron. Here *L_i_* (**z***_j_*) is the irradiance field of the *i*^th^ E-field at the location of the *j*^th^ neuron with absorption cross-section *σ* = 0.1 μm^2^ at full fluorophore activation.

With no beam steering, each E-field corresponds to its one E-pixel. But with beam steering, each E-pixel has several E-fields; we used 9 E-fields per E-pixel in all designs with steerable fields.

### Detection matrix

The detection matrix **D** (D-pixels × neurons ↦ probability) comprises elements

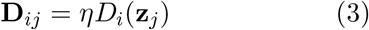

expressing the probability of a fluorescent photon emitted by the *j*^th^ neuron being detected by the *i*^th^ D-pixel. Here *D_i_*(**z***_j_*) is the detection field of the *i*^th^ D-pixel sampled at the location of the *j*^th^ neuron and *η* = 0.6 is the detector quantum efficiency.

### Illumination cycle

The illumination cycle matrix **J** (frames × E-fields) is an indicator matrix (consisting of 0s and 1s) specifying one full illumination cycle of E-field activations. Each row specifying an acquisition frame comprising the active E-fields. All the frames together comprise a single measured *sample* of duration Δ*t*. We used Δ*t* = 2 ms. However, the results of this study are insensitive to this setting as long as Δ*t* is short relative to the calcium dynamics.

The simplest illumination cycle would have a diagonal **J**, simply to cycling through each E-pixels’ individual illumination fields activating them one at a time. This cycle would deposit only *w_E_* = 100 μW of light and yield excessively large acquired data since the frames would need to be very short. Therefore, a pattern with many simultaneously active E-fields would potentially perform better.

We applied a simple greedy heuristic algorithm to produce more practical illumination cycles to reduce the number of frames. Starting with a diagonal **J**, we computed the *cross-exposure matrix* **Q** (frames × frames):

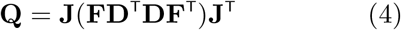

expressing the amount of overlap in detector exposure between pairs of frames. Finding the smallest element **Q***_ij_* revealed the two frames that had minimal overlap in D-pixel exposure. These frame pairs were combined by adding the corresponding rows of **J** together until achieving the desired rank.

The number of frames varied between 79 and 277 frames, depending on the design. Thus each frame was 7–25 μs in various designs.

### Mixing matrix

The optical signal mixing matrix **A** (channels × neurons ↦ photons) comprises elements

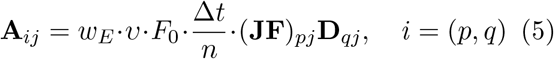

where the *i*^th^ channel combines the *q*^th^ D-pixel in the *p*^th^ frame of the illumination cycle. The emitter power is *w_E_* = 100 μW and the number of photons per joule is *ν* = 2.4 × 10^18^ (Table 1).

Thus the number of channels in a sample is the product of the number of frames (the rank of **J**) and the number of D-pixels.

Then given the vector of the neurons’ fluorescence states (as fractions of max fluorescence), the expected value of the signal (photon counts) picked up by the probe will be

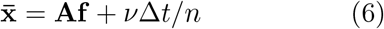

Here *v* = 300 (s ^−1^) is the dark photon count rate of the D-pixels, Δ*t*/*n* is the frame duration.

The acquired signal will be contaminated by quantal noise: uncorrelated variations with independent Poisson distribution with variances equal to 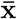. Thus the covariance matrix of the acquired signal will be

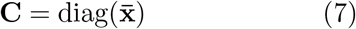

### Variance-normalized mixing matrix

Rather than carrying the dark counts and the covariance matrix through all the computations, we will subtract the dark current and, assuming that we can estimate **C** from the acquired signal, calculate the *noise-normalized mixing matrix*:

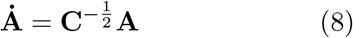

such that, if **C** is estimated for fluorescence vector **f**_0_, then for small deviations **f** = **f**_0_ + *δ*, the mean and variance of the acquired signals become

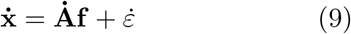

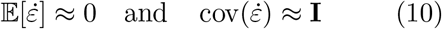

In the simulation, we used **f**_0_ = 0.05 · **1** (vector of constants corresponding to 5% average fluorescence) and *δ* was a vector of zeros everywhere except for the spiking neuron, where it had the value 0.015 corresponding to Δ*F/F*_0_ = 0.4.

### Demixing matrix

The demixing matrix **W** (neurons × channels) recovers an estimate of **f** as

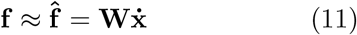

To stabilize the solution in the likely cases when 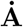 may be ill-conditioned, we use *L*_2_-regularized pseudo inverse [10], which solves the optimization problem finding the closest estimate of the fluorescence vector that explains the data but with a penalty term imposed on the magnitude (*L*_2_-norm) of the estimate.

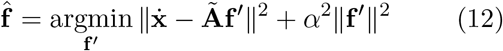

where *α* is the regularization parameter.

This optimum is attained with

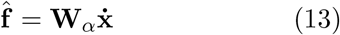

with the demixing matrix

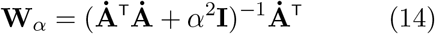

It may also be expressed in terms of the singular value decomposition 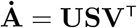:

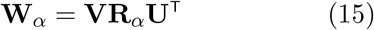

with

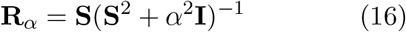

This effectively suppresses the effects of singular values that are not substantially higher than *α*.

If *S*_max_ is the largest singular value in **S**, then

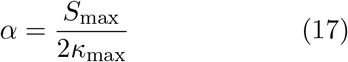

ensures that the condition number of the inverse (the ratio of the largest value of **R***_α_* to its lowest value) does not exceed *κ*_max_. We set *κ*_max_ = 10^6^ for numerical stability.

The regularized solution has the bias

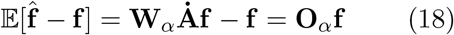

Then the Euclidean norms of the rows ∥*o_i_*∥ of the bias matrix

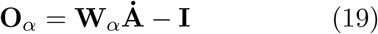

express the relative contamination of estimate of the signal from the *i*^th^ neuron by signals from other neurons. We considered cells whose relative bias was *o_i_* > 0.01 from SNR estimation. The regularization process “sacrifices” these cells to stabilize the rest of the solution.

### Spike SNR

For the majority of neurons that pass the bias criterion, the estimate 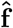 can be considered unbiased, so that

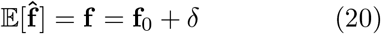

and the noise in the estimate can be obtained from the covariance matrix:

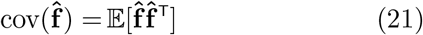

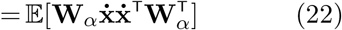

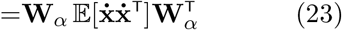

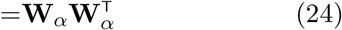

and

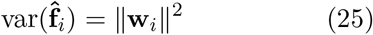

where ∥w*_i_*∥ is the Euclidean norm of the *i*^th^ column of **W***_α_*.

Thus the SNR for a small event of magnitude *δ_i_* in *i*^th^ neuron in one sample will be

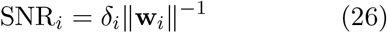

Shot noise is uncorrelated across samples and the SNR can be improved by applying a temporal filter. For example, if the calcium signal is modeled as an instant-rise/exponential-decay transient with max amplitude *δ* and time constant *τ*_Ca_ = 1500 ms, when the signal is sampled at 500 Hz (Δ*t* = 2 ms), then the matched filter *h*(*t*) will improve the SNR by the factor of

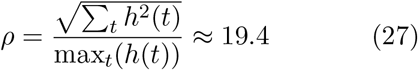

relative to the peak SNR in the unfiltered signal.

The SNR value shown in Eq. 1 and used in Figures 4 and 5 is the filtered spike SNR:

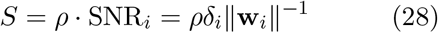

### SNR factorization

The noise-normalized matrix 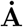 maps neuronal activity *δ* into the space of probe channels with spherical unity noise. This means that the recorded differences 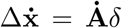 have mixed signal-to-noise ratios of 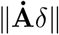. For an event of amplitude *δ_i_* isolated to the *i*^th^ neuron, this will translate into its mixed SNR of

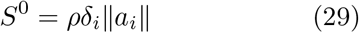

where ∥*a_i_*∥ is the Euclidean norm of the *i*^th^ column of 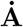.

The demixing matrix **W***_α_* is the regularized pseudoinverse of 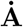, so that (with the exception of the bias introduced through regularization in a limited number of neurons),

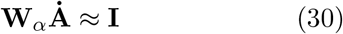

This means that the dot products between the *j*^th^ rows of **W***_α_* and the *i*^th^ columns of 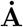 will be

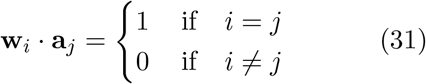

In other words, the vector **w***_i_* is orthogonal to the hyperplane spanned by the activity of all the other neurons—other than the *i*^th^ neuron—in the space of the recorded signal. Furthermore, by its construction, **w***_i_* is also the closest such vector to **a***_i_* considering in high dimensions, a hyperplane can have multiple norms. The cosine of the angle *θ* between these vectors is obtained by normalizing their dot product:

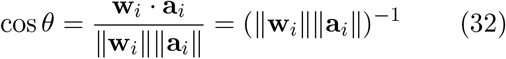

Finally, combining Eqs. 28, 29, and 32 we obtain the decomposition in Eq. 1.

## Discussion

In this study, we define a framework for modeling and evaluating the potential performance of various architectures of integrated neurophotonic systems. The simulations show that this new paradigm of compact and implantable lensless functional imagers can indeed provide dense coverage of circuit-level activity at arbitrary brain depths. The simulation also elucidate the subtle design tradeoffs involved in attaining optimal performance. With only a few iterations of design optimization, the study demonstrates significant improvements in the potential for separability of optical signals recorded with such probes. The framework now allows launching a systematic effort stign optimization under the constraints of manufacturability and minimal tissue damage. Continuing this work will also identify optical schemes for inferring the demixing matrix for real experiments as well as source localization.

## Acknowledgements

We thank Joseph Redford, a graduate student in Dr. Roukes’ lab, for implementing light transport simulations used for earlier versions of this analysis circa 2015.

